# An oscillating tragedy of the commons in replicator dynamics with game-environment feedback

**DOI:** 10.1101/043299

**Authors:** Joshua S. Weitz, Ceyhun Eksin, Keith Paarporn, Sam P. Brown, William C. Ratcliff

## Abstract

A tragedy of the commons occurs when individuals take actions to maximize their payoffs even as their combined payoff is less than the global maximum had the players coordinated. The originating example is that of over-grazing of common pasture lands. In game theoretic treatments of this example there is rarely consideration of how individual behavior subsequently modifies the commons and associated payoffs. Here, we generalize evolutionary game theory by proposing a class of replicator dynamics with feedback-evolving games in which environment-dependent payoffs and strategies coevolve. We initially apply our formulation to a system in which the payoffs favor unilateral defection and cooperation, given replete and depleted environments respectively. Using this approach we identify and characterize a new class of dynamics: an oscillatory tragedy of the commons in which the system cycles between deplete and replete environmental states and cooperation and defection behavior states. We generalize the approach to consider outcomes given all possible rational choices of individual behavior in the depleted state when defection is favored in the replete state. In so doing we find that incentivizing cooperation when others defect in the depleted state is necessary to avert the tragedy of the commons. In closing, we propose new directions for the study of control and influence in games in which individual actions exert a substantive effect on the environmental state.

---

Game theory is based on the principle that individuals make rational decisions regarding their choice of actions given suitable incentives [1, 2]. In practice, the incentives are represented as strategy-dependent payoffs. Evolutionary game theory extends game theoretic principles to model dynamic changes in the frequency of strategists [3]. Replicator dynamics is one commonly used framework for such models. In replicator dynamics, the frequencies of strategies change as a function of the social makeup of the community [4–6]. For example in a snowdrift game (also known as a hawk-dove game), individuals defect when cooperators are common but cooperate when cooperators are rare [2]. As a result, cooperation is predicted to be maintained amongst a fraction of the community [4, 6]. Whereas, in the prisoner’s dilemma individuals are incentivized to defect irrespective of the fraction of cooperators. This leads to domination by defectors [6, 7].

Here, we are interested in a different kind of evolutionary game in which individual action modifies both the social makeup and environmental context for subsequent actions. Strategy-dependent feedback occurs across scales from microbes to humans in public good games and in commons’ dilemmas [8–11]. Amongst microbes, feedback may arise due to fixation of inorganic nutrients given depleted organic nutrient availability [12, 13], the production of extracellular nutrient-scavenging enzymes like siderophores [14–16] or enzymes like invertase that hydrolyze diffusible products [17], and the release of extracellular antibiotic compounds [18]. The incentive for public good production changes as the production influences the environmental state. Such joint influence occurs in human systems, e.g., when individuals decide to vaccinate or not [19–21]. Decisions not to vaccinate have been linked most recently to outbreaks of otherwise preventable childhood infectious diseases in Northern California [22]. These outbreaks modify the subsequent incentives for vaccination. Such coupled feedback also arises in public goods dilemmas involving water or other resource use [23]. In a period of replete resources there is less incentive for restraint [24]. Yet, over-use in times of replete resource availability can lead to depletion of the resource and changes in incentives.

In this manuscript we propose a unified approach to analyze and understand feedback-evolving games (Figure 1). We term this approach “co-evolutionary game theory”, denoting the coupled evolution of strategies and environment. The key conceptual innovation is to extend replicator dynamics [4] to include dynamical changes in the environment. In that sense, our approach is complementary to recent efforts to consider the evolvability of payoffs in a fixed environment [25]. Here, changes in the environment modulate the payoffs. In so doing, we are able to address problems in which individual behavior constitutes a non-negligible component of the system. As a case study, we revisit the originating tragedy of the commons example [24] and ask: what happens if over-exploitation of a resource changes incentives for future action? As we show, the cumulative feedback of decisions can subsequently alter incentives leading to new dynamical phenomena and new challenges for control.

**FIG. 1:**
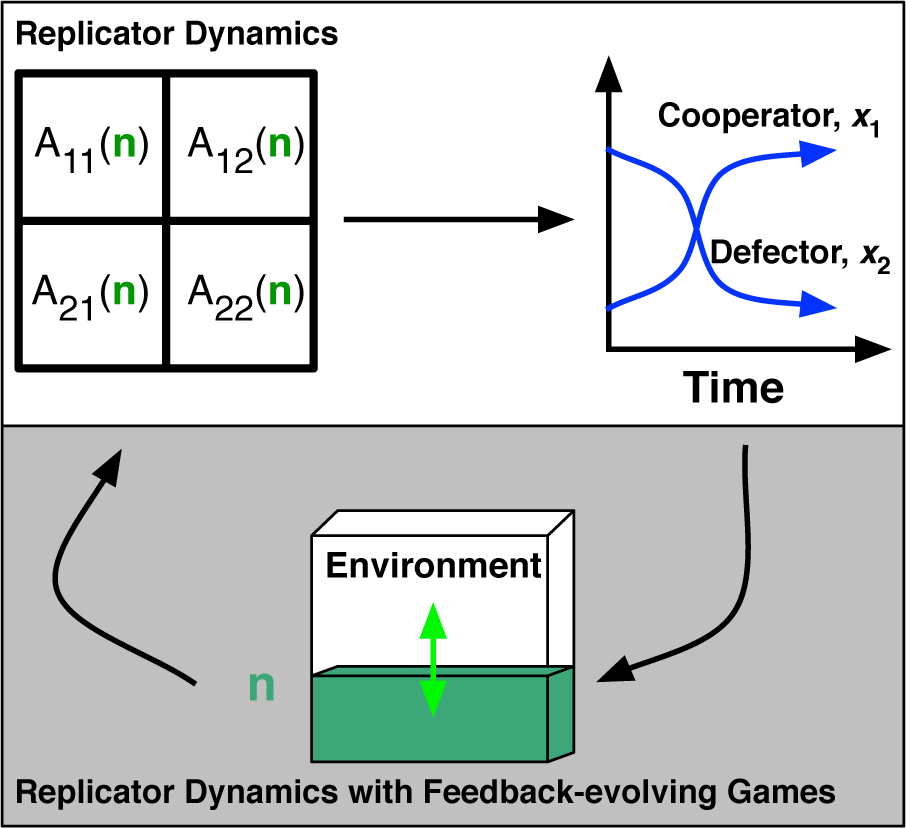
Schematic of replicator dynamics in feedback-evolving games. (Top) In replicator dynamics, the payoff matrix *A* determines frequency-dependent changes in strategies, *x_i_*. (Top and bottom) In replicator dynamics with feedback-evolving games, the frequencies of strategies influences the environment *n*, which, in term modifies the payoffs, *A*(*n*). The coupled system includes dynamics of both the payoff matrix and the strategies.

## Significance Statement

Classical game theory addresses how individuals make decisions given suitable incentives, e.g., whether to utilize a commons rapaciously or with restraint. Yet classical game theory does not typically address the consequences of individual actions that reshape the environment over the long-term. In this manuscript we propose a unified approach to analyze and understand the coupled evolution of strategies and the environment. We revisit the originating tragedy of the commons example and evaluate how over-use of a commons resource changes incentives for future action. In doing so, we identify an oscillatory tragedy of the commons in which the system cycles between deplete and replete environments and cooperation and defection behavior, highlighting new challenges for control and influence of feedback-evolving games.

## Methods and Results

### The context – evolutionary game theory as modeled via replicator dynamics

Here we introduce evolutionary game theory in the context of the prisoner’s dilemma as a means to motivate our co-evolutionary game theory formalism. Consider a symmetric two player game with strategies *C* and *D*, denoting cooperation and defection respectively. A standard instance of the payoff matrix is the prisoner’s dilemma (PD) in which the payoffs can be written as:

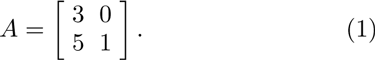

In this game, the player *C* receives a payoff of 3 and 0 when playing against player *C* and *D*, respectively. Similarly, the player *D* receives a payoff of 5 and 1 when playing against player *C* and *D*, respectively. These payoffs are commonly referred to as the reward for cooperation, *R* = 3, sucker’s payoff, *S* = 0, temptation to cheat, *T* = 5, and punishment for cheating, *P* = 1. Here, *R* < *T* and *S* < *P* so that mutual defection is the Nash equilibrium.

In evolutionary game theory, such payoffs can be coupled to the changes in population or strategy frequencies, *x*_1_ and *x*_2_, e.g., where *x*_1_ and *x*_2_ denote the frequency of cooperators and defectors such that *x*_1_ + *x*_2_ = 1. The coupling is expressed via replicator dynamics. The standard replicator dynamics for two-players games can be written as

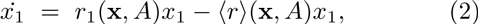

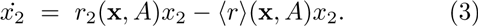

where *r*_1_, *r*_2_ and 〈*r*〉 denote the fitness of player 1, the fitness of player 2, and the average fitness respectively. In this convention then

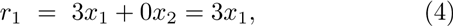

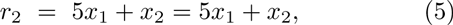

and the average fitness is:

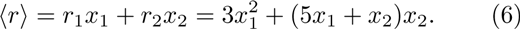

Because *x*_1_ + *x*_2_ = 1, we can rewrite the dynamics of *x* ≡ *x*_1_ as

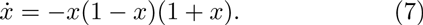

The replicator dynamics for the PD in Eq. (7) has 3 fixed points, but only two in the domain [0, 1], i.e., *x^*^* = 0 and *x^*^* = 1. The stability can be identified from the sign of the cubic, i.e., *x^*^* = 0 is stable and *x^*^* = 1 is unstable. This means that in the long-term a population with a minority of *D* players will, over time, change to one with a minority of *C* players, and the elimination of *C* players altogether.

In general, replicator dynamics for symmetric two-player games with a fixed payoff matrix can be written as:

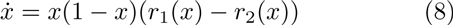

where the convention is again that *x* ≡ *x*_1_ and that *x*_1_ + *x*_2_ = 1. This formulation implies that the frequency of strategy 1 in the population will increase if the frequency-dependent payoff of strategy 1 exceeds that of strategy 2, and vice-versa. We can leverage this simple representation to consider the replicator dynamics given an alternative game:

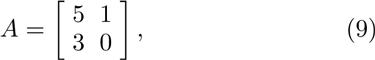

where *R* = 5, *S* = 1, *T* = 3, and *P* = 0. As is evident, *R* > *T* and *S* > *P* so that mutual cooperation is the Nash equilibrium. Here *r*_1_ = 4*x* + 1 and *r*_2_ = 3*x* such that

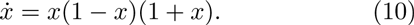

Again, the stability can be identified from the sign of the cubic, i.e., *x^*^* = 1 is stable and *x^*^* = 0 is unstable. The payoffs have changed such that cooperation is now a Nash equilibrium and a population with a minority of *C* players will, over time, change to one with a majority of *C* players, and eventually the elimination of *D* players.

### A model of replicator dynamics with feedback-evolving games

We consider a modified version of the standard replicator dynamics in which:

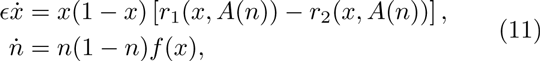

where *f*(*x*) denotes the feedback of strategists with the environment and the term *n*(1 − *n*) in Eq. (11) ensures that the environmental state is confined to the domain [0, 1]. The value of *ϵ* is a property of the agents and denotes the relative speed by which individual actions modify the environmental state. What distinguishes the model is that the payoff matrix *A*(*n*) is *environment-dependent* and that strategy and environmental dynamics are coupled (see Figure 1). The state of the environment is characterized by the scalar value, *n*. The environmental state changes as a result of the actions of strategists, such that the sign of *f*(*x*) denotes whether *n* will increase or decrease, corresponding to environmental degradation or enhancement when *f* < 0 or *f* > 0 respectively. Finally, the rate of environmental dynamics is set, in part, by the dimensionless quantity *ϵ*, such that when 0 < *ϵ* ≪ 1 then environmental change is relatively slow when compared to the change in the frequency of strategists.

Initially, we evaluate this class of feedback-evolving games via the use of the following environment-dependent payoff matrix:

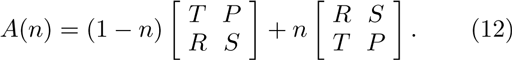

We retain the assumption of the prior section that *R* > *S* and *T* > *P*. As such, this initial class of games has an embedded symmetry such that mutual cooperation is a Nash equilibrium when *n* = 0 and mutual defection is a Nash equilibrium when *n* = 1. This state-dependent payoff matrix can be written as

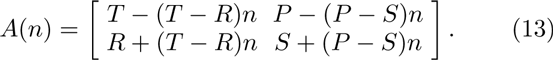

The payoff matrix *A*(*n*) interpolates between the two scenarios described in the previous section. Cooperation or defection are favored in the limits of *n* → 0 or *n* → 1, respectively. In addition, we assume that the environmental state is modified by actions of the population:

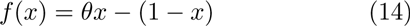

in which *θ* > 0 is the ratio of the enhancement rates to degradation rates of cooperators and defectors, respectively. The payoff-dependent fitnesses are

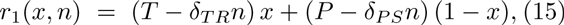

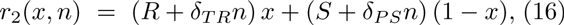

given *δ_PS_ = P − S* and *δ_TR_ = T − R*, such that the complete model can be written as:

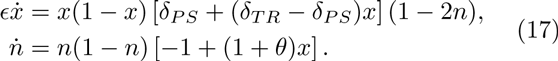

There are five fixed points of this model of replicator dynamics with feedback-evolving games. Of these, four represent “boundary” fixed points, that is (i) (*x^*^* = 0, *n^*^* = 0) - defectors in a degraded environment; (ii) (*x^*^* = 0, *n^*^* = 1) - defectors in a replete environment; (iii) (*x^*^* = 1, *n^*^* = 0) - cooperators in a degraded environment; and (iv) (*x^*^* = 1, *n^*^* = 1) - cooperators in a replete environment. There is also an interior fixed point, 
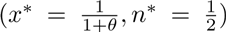
, representing a mixed population of cooperators and defectors in an intermediate environment. In Appendix A we prove that all of the boundary fixed points are unstable and the interior fixed point is neutrally stable. Further, we show that the system has a constant of motion. As a consequence, the global dynamics correspond to closed periodic orbits in the interior of the domain for any initial condition in which *x*_0_ ∈ (0, 1) and *n*_0_ ∈ (0, 1), with the exception of the interior fixed point. None of these orbits are limit cycles given the symmetries imposed in the payoff matrix.

The orientation of orbits in the phase plane defined by (*x, n*) should be counter-clockwise. The intuition is as follows. A system initialized in the interior near to (*x* = 0, *n* = 0) will rapidly move towards (*x* = 1, *n* = 0), as cooperation is favored. Then, as cooperators enhance the environment, the system will be driven closer to (*x* = 1, *n* = 1). Defectors will invade an environmentally enhanced state and the system will rapidly move to one near (*x* = 0, *n* = 1). Finally, in an environment dominated by defectors, the environmental state will be degraded and the system will be driven closer to (*x* = 0, *n* = 0).

This intuition holds throughout the domain. Dynamics of the system given the choice of payoffs, *R* = 3, *S* = 0, *T* = 5 and *P* = 1, and different initial conditions are shown in Figure 2.

**FIG. 2:**
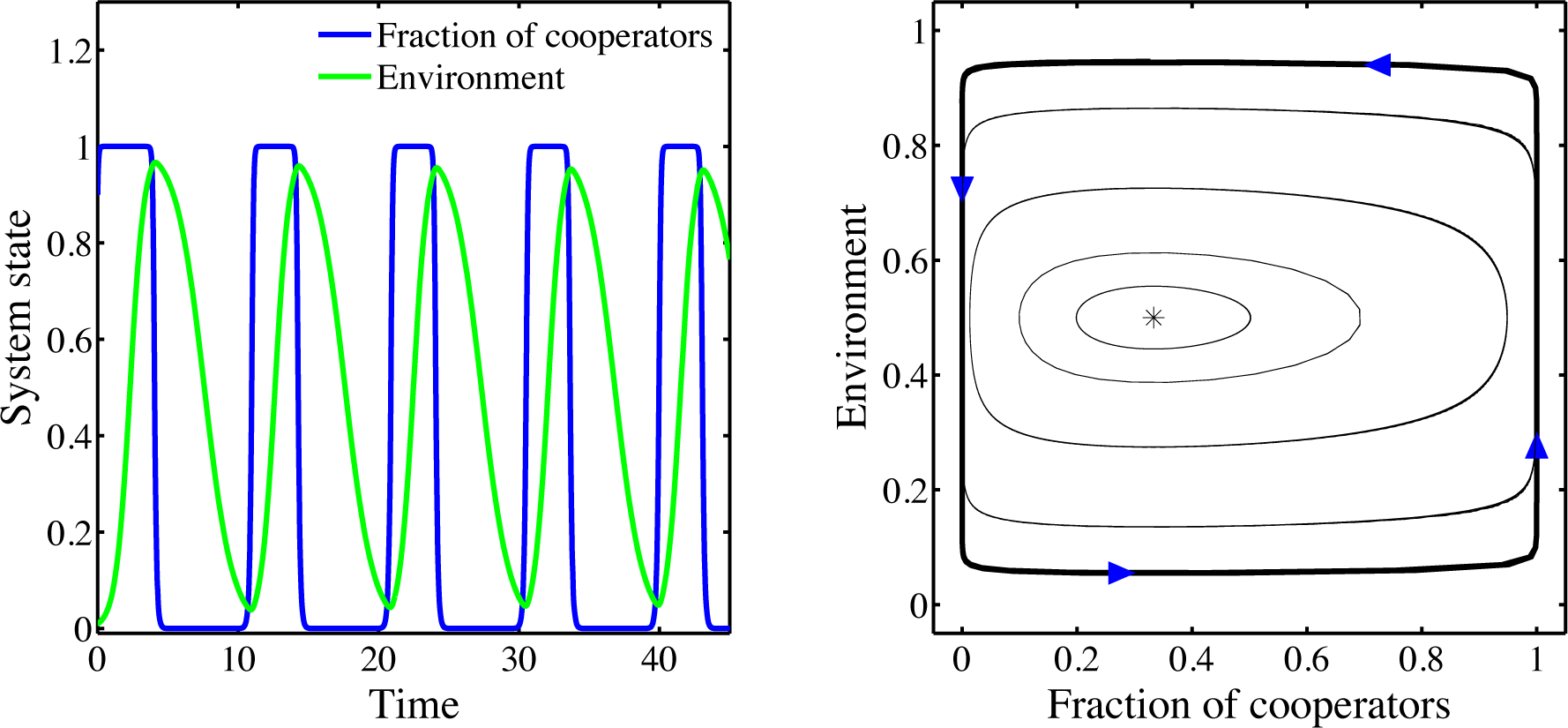
Persistent oscillations of strategies and the environment. (Left) Time series of the fraction of cooperators *x* (blue) and the environmental state *n* (green), correspond to dynamics arising from Eq. (17) with *ϵ* = 0.1, *θ* = 2, and the payoffs *R* = 3, *S* = 0, *T* = 5, and *P* = 1. (Right) Phase plane dynamics of *x − n* system. The arrows denote the direction of dynamics with time. The distinct curves correspond to initial conditions (0.9, 0.01), (0.8, 0.15), (0.7, 0.3), (0.5, 0.4), (0.4, 0.45). Appendix A includes a proof of the existence of a constant of motion associated with these dynamics. The asterisk denotes the predicted neutrally-stable fixed point at (1*/*3, 1*/*2).

### Generalized conditions for an oscillating tragedy of the commons

Here, we generalize our analysis by considering a model of replicator dynamics with feedback-evolving games with asymmetric payoffs:

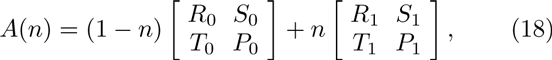

where 0 ≤ *n* ≤ 1. As before, we assume that if *n* = 0 then the payoff matrix has a unique Nash equilibrium corresponding to cooperative dominance, *i.e.,*, *R*_0_ > *T*_0_ and *S*_0_ > *P*_0_. Similarly, we assume that if *n* = 1 then the payoff matrix has a unique Nash equilibrium corresponding to defector dominance, *i.e., R*_1_ < *T*_1_ and *S*_1_ < *P*_1_. By breaking the symmetry of payoffs, we are able to explore cases in which both the relative sign and the magnitude of payoffs changes as a function of the environmental state.

As before, this system has five fixed points, of which four correspond to unstable fixed points on the boundary. In Appendix B we prove that the system has an unstable interior fixed point when

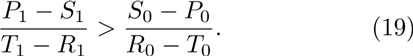

Here the qualitative outcomes depend on both the *sign* and *magnitude* of differences between payoffs, rather than the signs alone. Numerical simulations exhibit oscillations when Eq. (19) is satisfied (see Figure 3-top for an example). The numerical simulations also indicate that the oscillations grow in magnitude, possibly approaching the boundary. This observation raises a question: do the asymptotic dynamics converge to a limit cycle in the interior or to a heteroclinic cycle on the boundary?

**FIG. 3:**
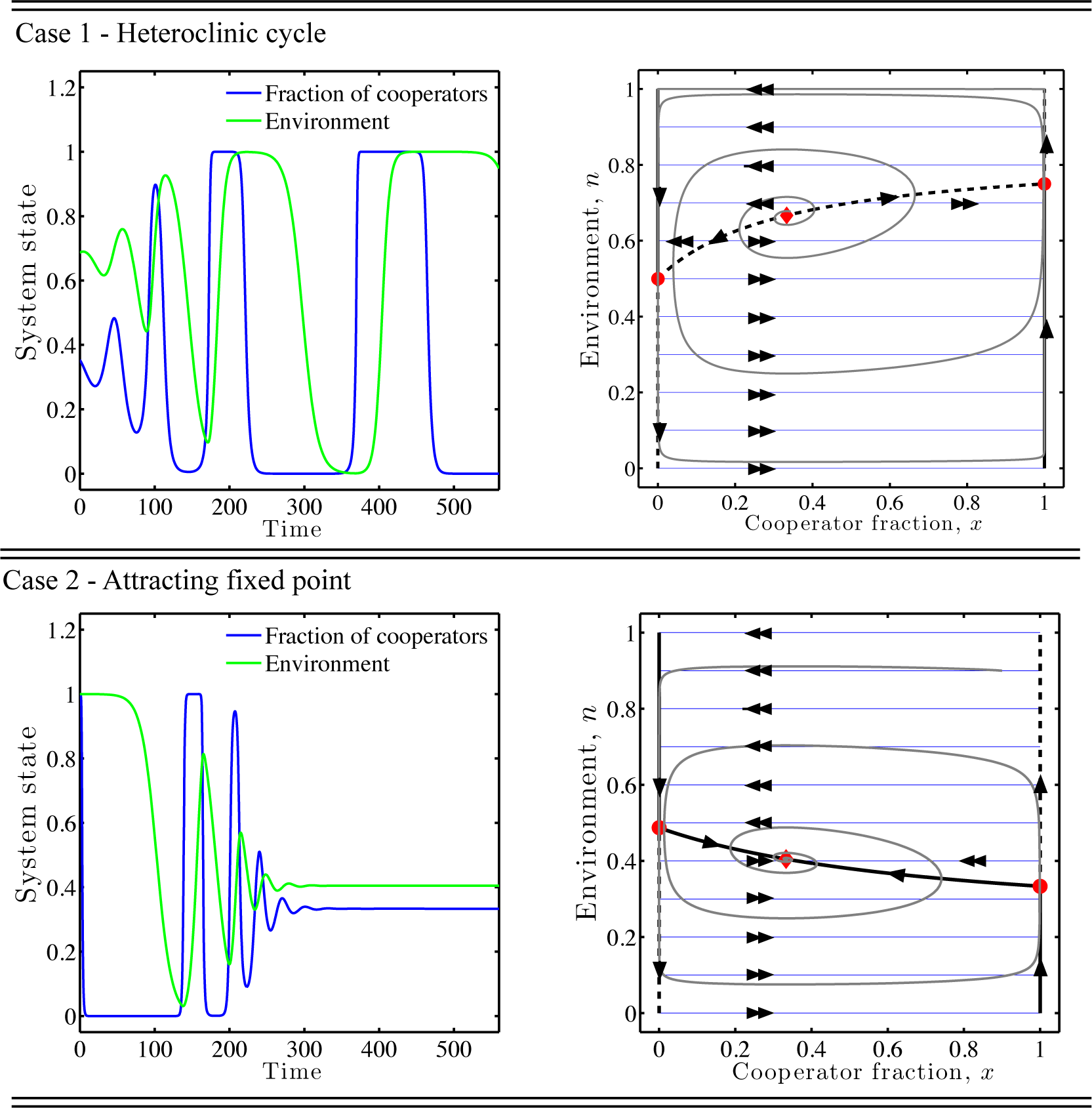
Fast-slow dynamics of feedback-evolving games, where *x* and *n* are the fast and slow variables respectively - including critical manifolds and realized dynamics. In both panels, the black lines denote the critical manifolds with solid denoting attractors and dashed denoting repellers. The blue lines and double arrows denote expected fast dynamics in the limit *ϵ* → 0. The red circles denote the bifurcation points of the fast subsystem parameterized by *n*. The single arrows denote expected slow dynamics. The gray curve denotes the realized orbit. In both cases, *ϵ* = 0.1 and *θ* = 2. (Top) Relaxation oscillations converging to a heteroclinic cycle arising due to a saddle-node bifurcation in the fast-subsystem parameterized by *n* in which the critical manifold is a repeller. The payoff matrix *A*(*n*) is that defined in Eq. (21). (Bottom) Relaxation oscillations converging to a fixed point arising due to a saddle-node bifurcation in the fast-subsystem parameterized by *n* in which the critical manifold is an attractor. The payoff matrix *A*(*n*) is that defined in Eq. (23).

In Appendix C we prove that the oscillations converge to an asymptotically stable, *heteroclinic cycle* and not to a limit cycle. We do so by leveraging conditions for the emergence of heteroclinic cycles in replicator dynamics [26, 27]. The heteroclinic cycle includes the four fixed points on the boundary in this order, (1, 1), (0, 1), (0, 0), (1, 0), which then return to (1, 1). The “tragedy” occurs given that dynamics initialized near (1, 1) are driven away to the environmental depleted state due to the actions of rational actors/agents acting in their own self-interest. Moreover, although it is not required for oscillations, we presume that *R*_0_ < *R*_1_, so that cooperation in the depleted state has less benefit than that of cooperation in the replete state. In summary, we use the term *Oscillating Tragedy of the Commons* to denote the emergence of oscillations that asymptotically connect the depleted and replete states. The use of “tragedy” has to do with the inevitability of environmental degradation (here, *n* = 0), or as Hardin emphasized “this inevitableness of destiny” [24].

The finding of an asymptotically stable heteroclinic cycle implies that dynamics initialized at an interior point and not located at a fixed point will approach the boundary. Over time, the dynamics will spend an ever increasing amount of time near a fixed point before “hopping” to the subsequent fixed point in the cycle. Near these fixed points the nonlinear dynamics are governed by linearized dynamics and two characteristic eigenvalues, one associated with an attractive “pull” towards the fixed point and one associated with a repelling “push” away from the fixed point. Hence, Eq. (19) can be interpreted as denoting the relative strength of the product of the stable (pull) vs. unstable (push) eigenvalues around the cycle (see Appendix C for details). Oscillations emerge when the pull towards the fixed points is stronger than the push away from the fixed points in a cycle. We also find that dynamics converge to an interior fixed point when Eq. (19) is not satisfied (see Figure 3-bottom for an example). In this event, the tragedy of the commons is averted and the system does not inevitably reach the depleted state.

### Fast-slow oscillatory dynamics in feedback-evolving games

In order to provide further intuition we leverage the fact that when 0 < *ϵ* ≪ 1 the dynamics correspond to that of fast-slow systems where *x* is the “fast” variable and *n* is the “slow” variable [28]. Later we will show that insights gained in the limit case hold irrespective of the relative rate change of environment and strategies. Consider a rescaling of time such that *τ* = *t/ϵ*, such that we rewrite Eq. (17) as:

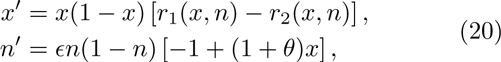

where the ′ denotes a derivative with respect to *τ*. For 0 < *ϵ* ≪ 1, this rescaling identifies *n* as the slow variable. Let 
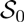
 denote the critical set of the system [28], i.e., the set of values of (*x, n*) for which *x′* = 0. The set 
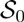
 is made up of multiple critical manifolds. So long as the system is not close to this critical set, then the dynamics of *x* are much faster than that of *n*, i.e., by a factor of order 1/*ϵ*. The critical manifolds of this system in the bounded domain 0 ≤ *x* ≤ 1 and 0 ≤ *n* ≤ 1 are: (i) *x* = 0; (ii) *x* = 1; and (iii) the set of points (*x_c_, n_c_*) that satisfy *r*_1_(*x_c_, n_c_*) = *r*_2_(*x_c_, n_c_*). The last of these critical manifolds can be interpreted as the interior nullcline of *x*. We assume that *n* parameterizes the dynamics of *x* far from the critical manifold.

As an example, consider the following payoff matrix:

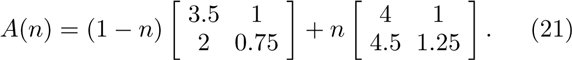

Given the payoff matrix in Eq. (21), the one-dimensional fast-subsystem can be written as:

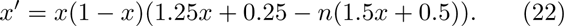

In this case, the interior critical manifold satisfies 
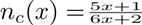
. For *n* > 3*/*4, there are two fixed points, *x* = 0 and *x* = 1, which are stable and unstable respectively. For *n* < 1*/*2, there are also two fixed points, *x* = 0 and *x* = 1, which are unstable and stable respectively. This system undergoes two saddle-node bifurcations at the values *n* = 1*/*2 and *n* = 3*/*4. For values of the slow variable 1*/*2 < *n* < 3*/*4, the system has three fixed points, such that *x* = 0 and *x* = 1 are stable and *x_c_* is unstable where 
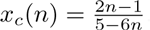
.

System dynamics can be understood in terms of a series of fast and slow changes. Consider initializing the system at values (*x*_0_, *n*_0_) with *n*_0_ < 1*/*2, i.e., in the region where there are only two fixed points of the fast-dynamics. The system will behave akin to a one-dimensional system and increase rapidly in *x*, parameterized by the value *n = n*_0_. As the system approaches the attracting critical manifold, *x* = 1, then the system dynamics will be governed by the slow variable dynamics, *n′*. Cooperators will enhance the environmental state, given that *n′* > 0 for *x* → 1. The system will then *slowly* approach the fixed point (1, 1). This fixed point is unstable in the fast direction, such that the system will *rapidly* approach the attracting critical manifold of *x* = 0, i.e., the point (0, 1). Again, the system will then *slowly* change in environmental state towards the point (0, 0), given that *n′* < 0 for *x* → 0. Now that *n* < 1*/*2, the system will be dominated by the fast subsystem dynamics, *rapidly* increasing *x* - completing the cycle. The resulting dynamics will appear as relaxation oscillations with slow changes in environment alternating with rapid changes in the fraction of cooperators. The dynamics overlaid with the critical manifolds for this system are shown in Figure 3-top. We find that the system dynamics will asymptotically approach a heteroclinic cycle when *n_c_*(0) < *n_c_*(1). The condition for this asymptotic behavior corresponds to that of Eq. (19) (see Appendix C and D).

The key to the emergence of relaxation oscillations is that the interior critical manifold is a repeller. This is not universally the case. A counter-example is when Eq. (19) is not satisfied, e.g.,:

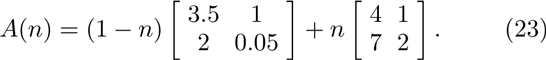

In this example, the overall dynamics converge to an asymptotically stable interior fixed point. The overall dynamics are again characterized by a mix of fast and slow changes, however they spiral in to the interior fixed point rather than away from it. The dynamics overlaid with the critical manifolds for this system are shown in Figure 3-bottom. We also find that the qualitative outcomes do not depend on the speed of the feedback (see Figure 4 and Appendix B and C for proof). Here, the invariance arises because of the separability of the dynamics so that the stability of the system is unaffected by the speed. This *ϵ*-invariance of qualitative outcomes is not universally the case for fast-slow systems [28].

**FIG. 4:**
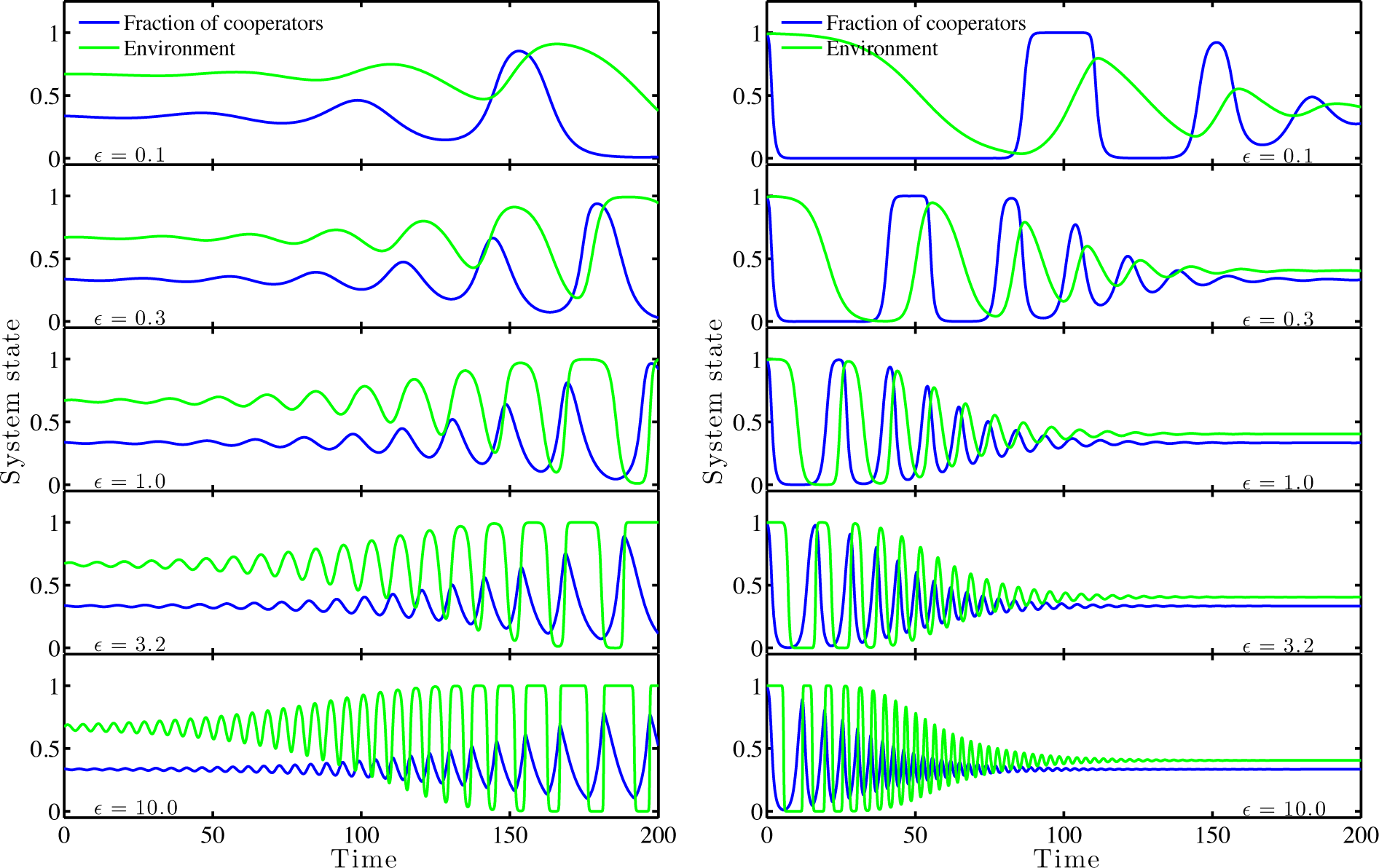
Invariance of system dynamics given change in the relative speed of strategy and environmental dynamics. The parameter *ϵ* is varied from 0.1 to 10 given cases where dynamics are expected to lead to a heteroclinic cycle (left) and to an interior fixed point (right). Other parameters are the same as in Figure 3. Although the transient dynamics differ, the qualitative dynamics remain invariant with respect to changes in *ϵ*.

### Generalized conditions for mitigating the tragedy of the commons given feedback-evolving games

The previous section identified conditions under which the tragedy of the commons is averted. In particular, the system converges to an intermediate environmental state when the cumulative strength of unstable eigenvalues around the cycle exceeds that of the stable eigen-values (Eq. (19)). This condition requires that mutual cooperation is a unique Nash equilibrium in a depleted state. Here, we ask: are there any other conditions in which a tragedy of the commons could be averted? To do so, we continue to fix the payoff structure of *A*_1_ to have a Nash equilibrium corresponding to mutual defection so as to analyze the effect of feedback on the tragedy of the commons. However in this section we consider *any ordering* of payoffs in *A*_0_ and analyze the corresponding replicator dynamics with feedback-evolving games.

There are four cases to consider corresponding to different combinations among the relative values of *T*_0_ and *R*_0_ as well *S*_0_ and *P*_0_. First, *A*_0_ may correspond to an anti-coordination game, i.e., *R*_0_ < *T*_0_, *S*_0_ > *P*_0_. Second, *A*_0_ may correspond to a coordination game, i.e., *R*_0_ > *T*_0_, *S*_0_ < *P*_0_. Third, *A*_0_ may have a unique Nash equilibrium corresponding to mutual cooperation, *i.e., R*_0_ > *T*_0_, *S*_0_ > *P*_0_. Fourth, *A*_0_ may have a unique Nash equilibrium corresponding to mutual defection, *i.e.,R*_0_ < *T*_0_, *S*_0_ < *P*_0_. Of these, we have already analyzed dynamics arising in the third case when *R*_0_ > *T*_0_, *S*_0_ > *P*_0_ (see Figures 2-4). The possible outcomes of dynamics that do not begin at a fixed point include convergence to a stable interior fixed point or persistent oscillations in the interior. The fourth case corresponds to domination by a defector strategy when *n* = 0 and when *n* = 1. Therefore, defection will be the dominant strategy for all values of *n*. The population will converge to *x^*^* = 0 and, by extension, to the depleted environmental state *n* = 0. There are no additional dynamics possible given the feedback structure studied here.

In Appendix D we find the fixed points and local stability for all values of payoffs of *A*_0_ in the two remaining cases. In the event that *A*_0_ constitutes an anti-coordination game, the system can converge to the boundary or to the interior. The stable boundary point corresponds to (*x_m_,* 0), where *x_m_* is the mixed Nash equilibrium of *A*_0_. Hence, the system exhibits a tragedy of the commons despite the fact that there is a mix of cooperators and defectors. The positive feedback of the cooperators is insufficient to improve the environment over the long-term. However, when the cooperator advantage asymmetry is sufficiently large, then the system converges to an interior fixed point. Counter-intuitively, this interior fixed point exhibits less cooperation than on the boundary. In the event that *A*_0_ constitutes a coordination game, the system converges to the stable fixed point (0, 0) corresponding to a tragedy of the commons, irrespective of initial conditions. The coupling between behavior and environment provides an “escape route” that leads to total defection in the system. To understand why, consider dynamics that ensue near the point (1, 0). In that event the payoff conditions favor cooperation, which drive the system closer to (1, 1), at which point the behavior switches to that of defection, i.e., approaching (0, 1). Defectors degrade the environment, converging to the locally stable fixed point (0, 0).

We summarize all possible dynamics in terms of a phase plane in Figure 5. The key point is that the tragedy of the commons can be averted when there is feedback between strategy and the environment. Convergence to an intermediate environmental state depends on the magnitude of payoffs in depleted environments as well as the relative strength at which cooperators enhance the environment. In this model, incentivizing the payoff to cooperate when others defect in a bad environment can help avert the tragedy of the commons.

**FIG. 5:**
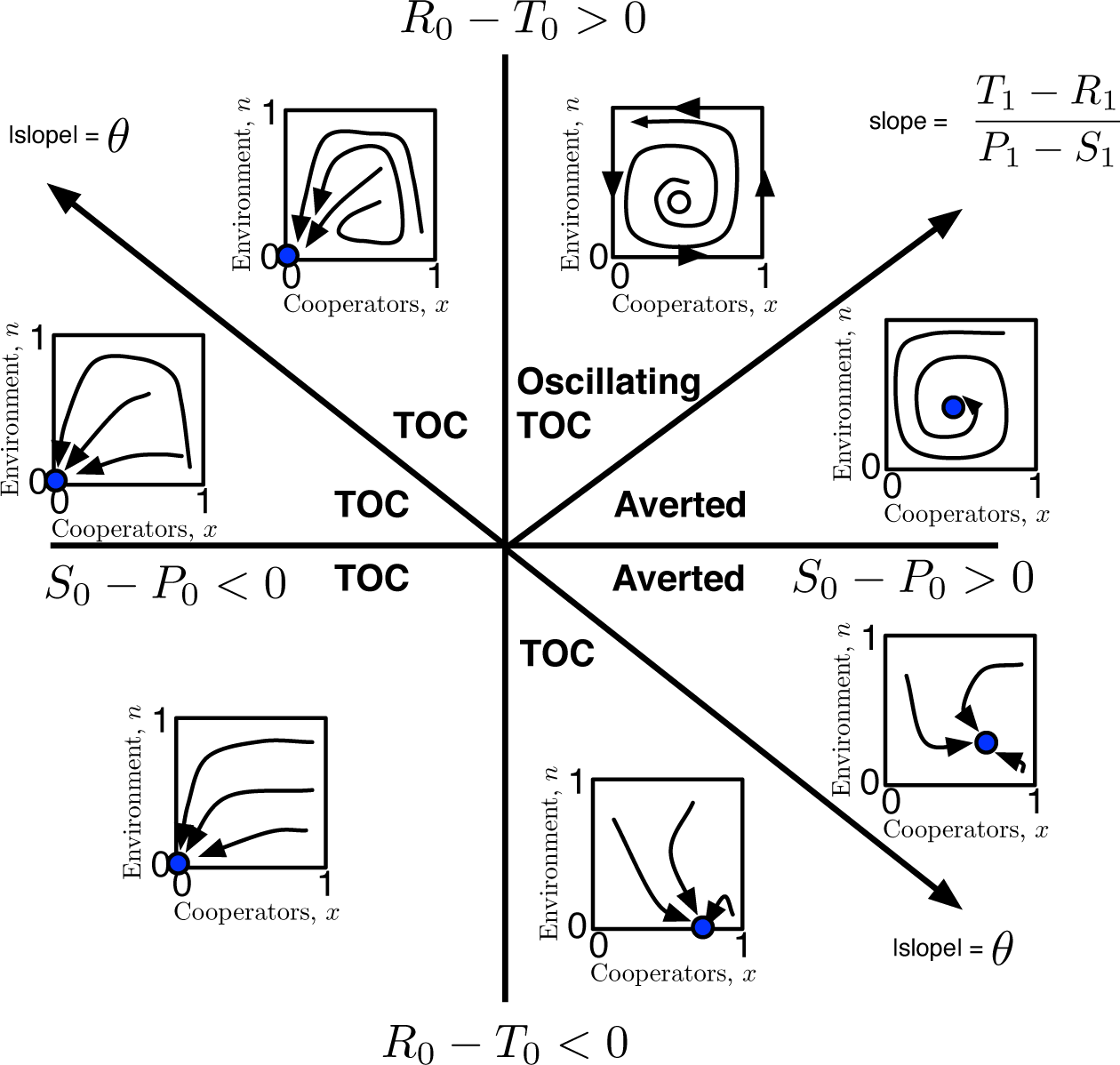
Summary of dynamics given all possible combinations of payoffs in the depleted state. The axes correspond to the relative values of *S*_0_ − *P*_0_ and *R*_0_ − *T*_0_. There are seven regions identified. The inset include schematics of dynamics corresponding to each combination of payoff values, where the closed circle denotes a locally stable fixed point. In all cases we consider a scenario in which the system has a unique Nash equilibrium corresponding to mutual defection in the replete state, *n* = 1. Appendix D includes detailed analysis and numerical simulations for the bottom-right and upper-left quadrants.

## Discussion

We proposed a co-evolutionary game theory that incorporates the feedback between game and environment and between environment and game. In so doing, we extended replicator dynamics to include feedback-evolving games. This extension is facilitated by assuming the environmental state can be represented as a linear combination of two different payoff matrices. Motivated by the study of the tragedy of the commons in evolutionary biology [29] we demonstrated how new kinds of dynamics can arise when cooperators dominate in deplete environments and when defectors dominate in replete environments. In essence, cooperators improve the environment, leading to a change in incentives that shift the game theoretic strategy towards defection. Repeated defections degrade the environment which re-incentives the emergence of cooperators. In this way there is the potential for a sustained cycle in strategy and environmental state, i.e., an oscillating tragedy of the commons (see Figures 3-4). Whether or not the cycle dies out or is persistent depends on the *magnitude* of payoffs. We also identified conditions under which a tragedy of the commons can be averted (see Figure 5).

Our proposed replicator model with feedback-evolving games considers the consequences of repetition in which the repeated actions of the game influences the environment in which the game is played. Thus, the model is complementary to long-standing interest in a different kind of repeated games, most famously the iterated prisoner’s dilemma [7, 30–35]. In such iterated games, strategies that include cooperation emerge, even if cooperation is otherwise a losing strategy in single-stage or one-shot versions of the game. Here, individuals do not play against another repeatedly or, posed alternatively, do not “recall” playing against another repeatedly. Instead, a feedback-evolving game changes with time as a direct result of the accumulated actions of the populations.

The feedback-evolving game analyzed here is closest in intent to a prior analysis of coupled strategy and environmental change in the context of durable public goods games [36]. That model assumed that fitness differences between producers and non-producers had no frequency-dependence and the environmental dynamics had at most a single fixed point. Unlike the present case, the model in [36] did not exhibit persistent oscillations. Here, the long-term dynamics depend on the magnitudes of payoffs, i.e., including both costs, benefits, and frequencies of alternate strategies, as well as the strength of feedback. For example, bacteria that produce a costly public good, i.e., cooperators, may have a selective advantage in a depleted environment when public goods are scarce. Cooperating bacteria can restore the availability of the public good, e.g., fixed nitrogen or excreted enzymes, thereby favoring defectors that consume but do not produce the public good. The emergence of defectors can, with time, degrade the environment.

We have shown (in Figures 2-4) that repeated oscillations of strategies and environmental state can arise when cooperation is favored in the depleted state. We have also classified the behavior of the model given all possible payoff matrices in the depleted state (see Figure 5 and Appendix D). In all other instances we find that global dynamics converge to a fixed point. This fixed point can be in the interior, i.e., corresponding to averting the tragedy of the commons. Averting the tragedy of the commons is possible, though not guaranteed, so long as cooperation is favored when others defect in the depleted state. The conditions for averting the tragedy of the commons in this model depends on the strength but not the speed of coupling. Alternative forms of feedback between strategy and environment [37, 38] as well as nonlinear combinations of payoff matrices may lead to novel dynamics. Density-dependent interactions may also lead to novel effects of social behavior on total population densities, not just their frequencies [39].

Thus far, we have assumed that the environment can recover from a nearly deplete state. The rate of renewal was assumed to be proportional to the cooperator fraction. In that sense, our work also points to new opportunities for control - whether for renewable or finite resources. Is it more effective to influence the strategists, the state, and/or the feedback between strategists and state? Analysis of feedback-evolving games could also have implications for theories of human population growth [40], ecological niche construction [41], and the evolution of strategies in public good games [25]. The extension of the current model to microbial and human social systems may deepen understanding of the short‐ and long-term consequences of individual actions in a changing and changeable environment [42]. We are hopeful that recognition and analysis of the feedback between game and environment can help to more effectively manage and restore endangered commons.

## Acknowledgments

This work was supported by a grant from the Army Research Office W911NF-14-1-0402 to JSW. JSW thanks Joel Cohen for feedback on an early draft of the manuscript and the organizers of the 2014 NAKFI workshop on Collective Behaviors: From Cells to Societies where work on this project began. The authors thank Michael Cortez, Jeff Shamma, and two anonymous reviewers for their comments on the manuscript, particularly the suggestion of one reviewer to investigate the asymptotic nature of oscillations in this model.

## Appendix A: Closed periodic orbits given replicator dynamics with feedback evolving games

In this section we show that replicator dynamics with feedback evolving games will exhibit closed period orbits given suitable symmetries in the payoffs in the good (*n* = 1) and bad (*n* = 0) environmental states. Consider an environment-dependent payoff matrix of the form:

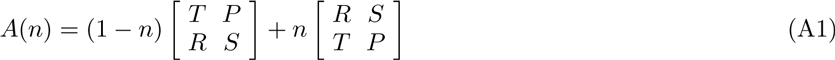

where *T* > *R* and *P* > *S*. In this case, mutual defection is the Nash equilibrium when *n* = 1 and mutual cooperation is the Nash equilibrium when *n* = 0. This payoff matrix can be rewritten as:

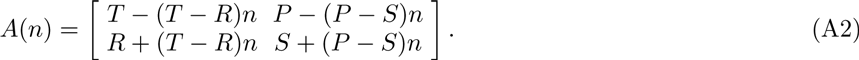

The dynamical system of equations are:

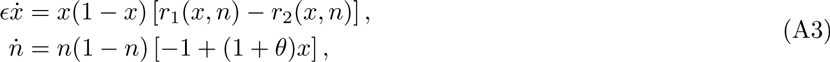

where

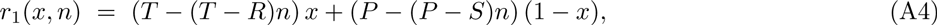

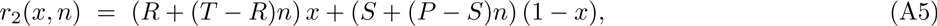

so that

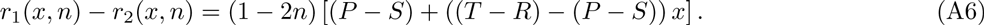

For convenience we denote the differences as *δ_PS_ = P* − *S* and *δ_T R_ = T* − *R*. In this notation, the system is:

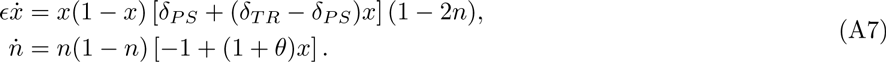

This system has five fixed points, including all four corners of the domain [0, 1]^2^ and the interior fixed point 
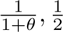
. Without loss of generality, we set *ϵ* = 1 to analyze the local stability of these fixed points. The Jacobian of the system at each of the four corners is:

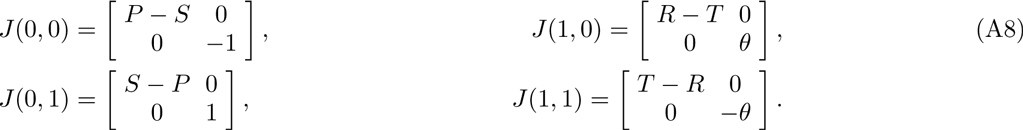

Because *T* > *R* and *P* > *S*, then each of these Jacobian has one positive eigenvalue and is locally unstable. In addition, the Jacobian of the interior fixed point is:

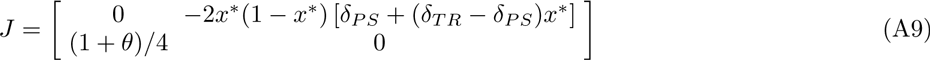

where 0 < *x^*^* = 1/(1 + *θ*) < 1 for the interior fixed point. The eigenvalues for the interior fixed point have no real components and the interior fixed point is neutrally stable.

We now analyze the global dynamics of the system and confirm that the intuition gained from analysis at the interior fixed point holds for the entire domain. The separability of the factors allows us to write

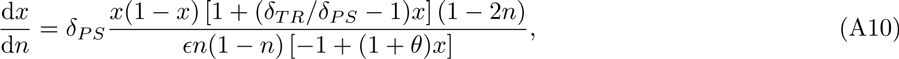

such that

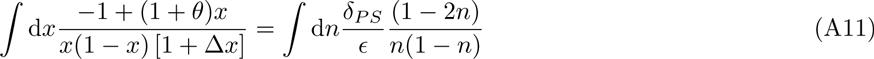

where 
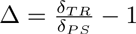
. The integral of the right-hand side of Eq. (A11) is

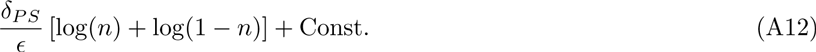

The integral of the left-hand side of Eq. (A11) is

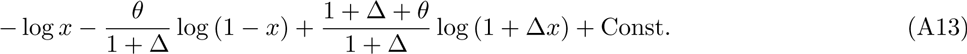

In this way, we identify the constant of motion

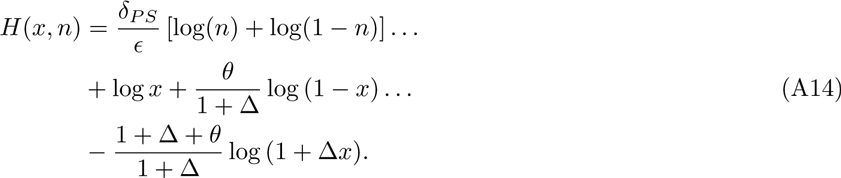

*H* is a constant of motion since

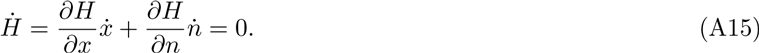

In summary, a system initialized with (*x*_0_, *n*_0_) in the interior of the square 0 < *x, y* < 1 and not at a fixed point will exhibit closed, periodic orbits.

By way of illustration, consider the example introduced in the main text:

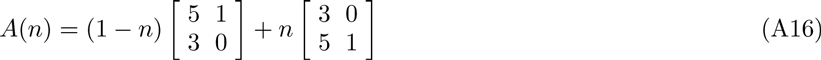

or, alternatively

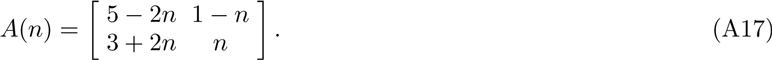

For this particular case, *δ_PS_* = 1 and Δ = 1, such that

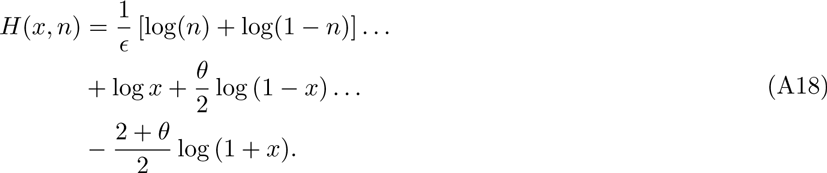

The constants of motion for each initial condition (0.9, 0.01), (0.8, 0.15), (0.7, 0.3), (0.5, 0.4), (0.4, 0.45) in Figure 2 are -49.8, -23.6, -18.2, -16.5, and -16.1, to three significant digits respectively.

## Appendix B: Stability analysis of replicator dynamics with feedback-evolving games

In this section we derive conditions for an oscillating tragedy of the commons in replicator dynamics with feedback-evolving games. Here, we consider the generalized payoff matrix

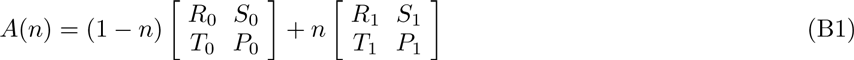

where 0 ≤ *n* ≤ 1, *R*_0_ > *T*_0_, *S*_0_ > *P*_0_, *R*_1_ < *T*_1_, and *S*_1_ < *P*_1_. Following the convention of Eq. 20, the dynamics are

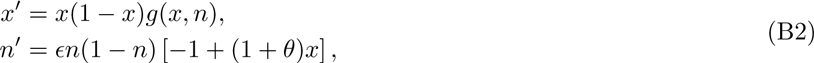

where *g*(*x, n*) = *r*_1_(*x*, *n*) − *r*_2_(*x, n*) denotes the differences in state-dependent payoffs and the ′ denotes a derivative with respect to *τ*. The interior fixed point (*x^*^, n^*^*) has an associated Jacobian:

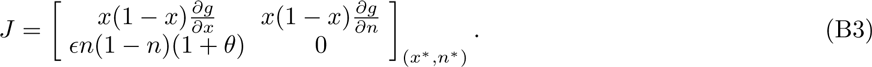

Here *∂g*/*∂n* < 0 for all cases of concern here because the replicator dynamics favor decreases in cooperation as an increasing function of the environmental state *n*. As such, the determinant is always positive. The stability of the fixed point depends on the sign of the trace of *J*. Because 0 < *x^*^* < 1, then the sign of the trace is equivalent to the sign of *∂g*/*∂x*. This result can also be anticipated by a fast-slow systems analysis in which the stability of any point on the nullcline depends strictly on *∂g*/*∂x*. Another consequence is that the trace does not depend on *ϵ*. Therefore, the qualitative dynamics will be the same for all values of *ϵ*, i.e., irrespective of the relative speed of the fast and slow variables (see Figure 4). Such an *ϵ*-invariance of qualitative phenomena is not universally the case in fast-slow dynamics.

Formally, *∂g*/*∂x* can be written as:

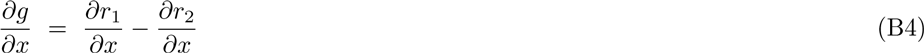

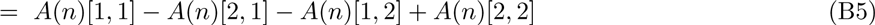

given the condition that *r*_1_ = *r*_2_ at a fixed point. We must solve for *n^*^* as a function of the payoff coefficients. As such, when *r*_1_ = *r*_2_ given *x = x^*^* = 1/(1 + *θ*), the following condition must be satisfied:

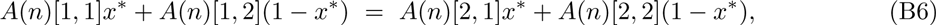

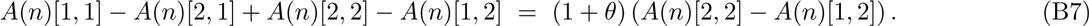

Recall that

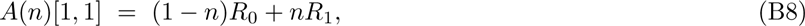

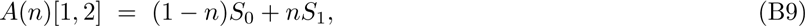

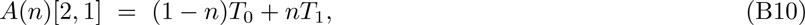

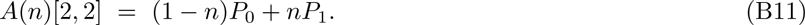

so that the condition for *n^*^* is

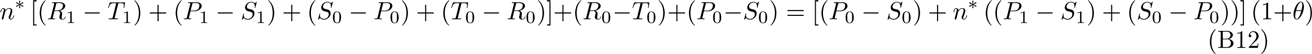

or equivalently,

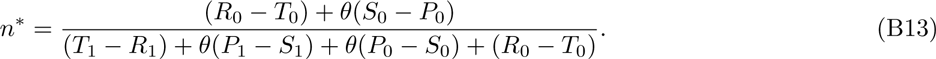

The point of interest *n^*^* satisfies *r*_1_ = *r*_2_. Substitution of terms at the fixed point yields

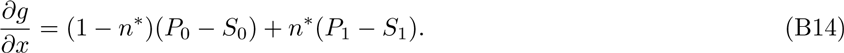

In other words the interior fixed point is unstable when *∂g/∂x* > 0, equivalently:

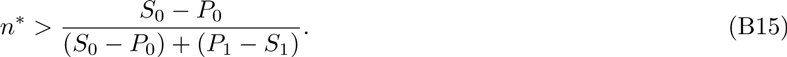

Recall that *S*_0_ > *P*_0_ and *P*_1_ > *S*_1_ in the games we consider and further that the r.h.s. of Eq. (B16) is equivalent to *n_c_*(0) in the main text, i.e., the intersection of the nullcline of *x́* with the (*x* = 0, *n*) boundary. We can then write the condition on instability as

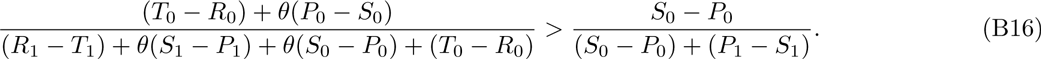

We denote the strictly positive dummy variables *C*_1_ = *T*_0_ − *R*_0_, *C*_2_ = *R*_1_ − *T*_1_, *D*_1_ = *P*_0_ − *S*_0_ and *D*_2_ = *S*_1_ − *P*_1_ and rewrite Eq. (B17) as:

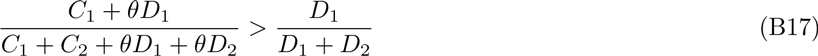

which after some algebraic re-arrangement yields

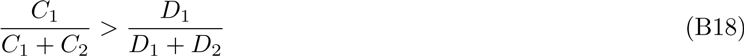

where we recognize the l.h.s. as

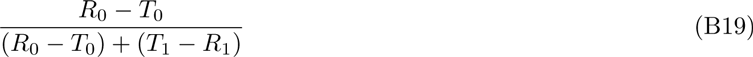

or equivalent to *n_c_*(1) in the main text, i.e., the intersection of the nullcline of *x́* with the (*x* = 1, *n*) boundary. To summarize, the interior fixed point is unstable when

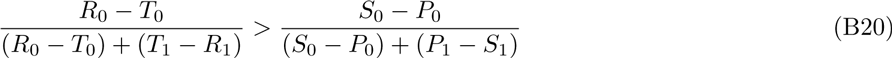

This can be further reduced to the Eq. (19) in the main text:

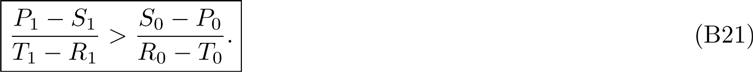

This condition is equivalent to the requirement that *n_c_*(0) < *n_c_*(1). We have already shown that all boundary fixed points are unstable. The only other rest point in the interior of the domain is unstable. In the next section we demonstrate that dynamics in the interior converge to an asymptotically stable *heteroclinic cycle*, i.e., attracting all points that are not initially at a fixed point to the boundary.

## Appendix C: Stability proof of heteroclinic cycle

We consider the planar dynamical system

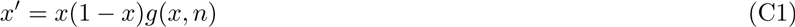

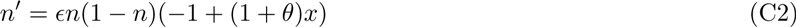

where

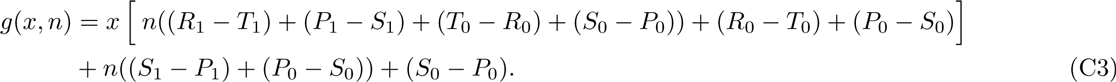

The state space of the system is the unit square *X* = [0, 1]^2^ with boundary *∂X* (the set of points on the edges), which is invariant under the dynamics. Using the condition in Eq. (19) of the main text, we prove the dynamical system has an attracting heteroclinic cycle Λ = *∂X* with orientation (0, 0) → (1, 0) → (1, 1) → (0, 1) → (0, 0) connecting the four corner fixed points along the boundary of the unit square. Under this condition, the four corner fixed points are saddles and the interior fixed point is unstable. The proof relies on properties of the characteristic matrix *C* of the heteroclinic cycle Λ (see Section 2, [1]), which encodes its stability properties and is constructed as follows. Observe that *X* is the intersection of four halfspaces,

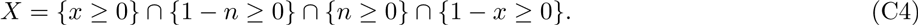

Define *x*_1_ = *x*, *x*_2_ = 1 − *n*, *x*_3_ = *n*, *x*_4_ = 1 − *x*. We can write the augmented dynamics as

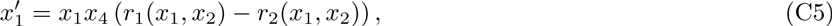

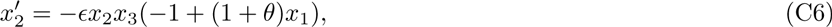

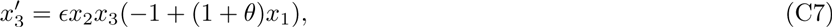

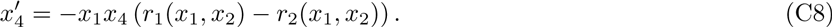

The characteristic matrix *C* has elements defined by

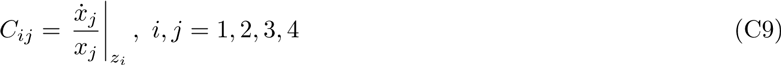

where *z_i_* is one of the four corner fixed points. We obtain *C* by following the scheme above:

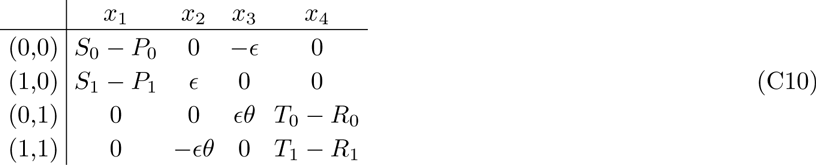

Recall that *S*_0_ − *P*_0_ > 0, *S*_1_ − *P*_1_ < 0, *T*_0_ − *R*_0_ < 0, and *T*_1_ − *R*_1_ > 0. We find that *C* has positive entries occurring only on the diagonal and each row and column contains only one positive entry. Thus, Λ is a *simple heteroclinic cycle* [1]. We review some technical conditions for the stability of heteroclinic cycles. From section 4 in [1], it is required that

- The heteroclinic cycle Λ is a compact invariant subset of the boundary *∂X*.
- Let Λ*_k_* ⊂ Λ for *k* = 1, …, *m* (*m* is the number of fixed points in Λ) be such that for each *x* ∈ Λ, there is a *k* with *ω*(*x*) ⊂ Λ*_k_* (*ω*(*x*) is the *ω*-limit set of *x*).

The above conditions are satisfied in our case since Λ = *∂X* is compact, *∂X* is invariant under the dynamics, and the four corner fixed points serve as the Λ*_k_*’s. The stability of Λ invokes the following result from [1]:

#### Theorem C.1

(Corollary 1 in [1]). Let Λ be a simple heteroclinic cycle, which is asymptotically stable within *∂X*. (This is automatically satisfied if the cycle is robust and all Λ*_k_* are fixed points.) Then *C* is a square matrix (after elimination of superfluous columns) with positive entries occurring only in the main diagonal (after a suitable rearrangement of the rows or columns). Let det *C* ≠ 0. If *C* is not an *M*-matrix (at least one leading principal minor is negative) then Λ is asymptotically stable.

Indeed, *C* is not an *M*-matrix by the following calculation and invoking Eq. (19) of the main text:

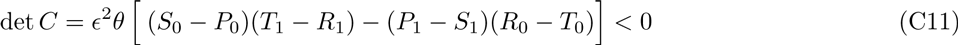

We note that this condition is equivalent to the interpretation from [2] that a robust heteroclinic cycle is asymptotically stable when the cumulative strength of the stable eigenvalues in the cycle exceeds that of the unstable eigenvalues. In addition we note that the condition is *ϵ*-invariant.

## Appendix D: Generalized dynamics of replicator dynamics with feedback evolving games

In this section we analyze the structure of fixed points and their local stability given variations in the state-dependent payoff matrix for replicator dynamics with feedback evolving games. As in the main text, we consider the symmetric 2 × 2 game defined for *n* ∈ [0, 1] by the matrix

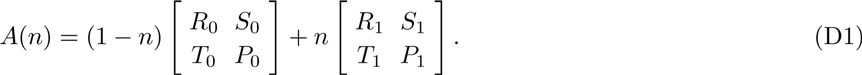

We are interested in the dynamics that occur for any combination of these eight parameter values given the restriction that *R*_1_ < *T*_1_, *S*_1_ < *P*_1_. Thus when *n* = 1, mutual defection is the Nash equilibrium. The replicator dynamics with feedback evolving games given *ϵ* = 1 are:

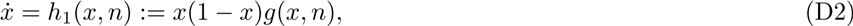

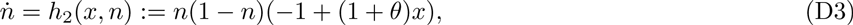

where *θ* > 0 and

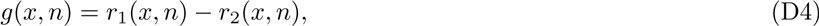

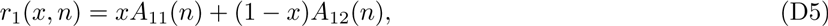

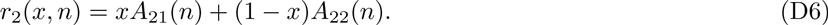

The four corner points are fixed points. Under certain conditions, there is an interior fixed point given by

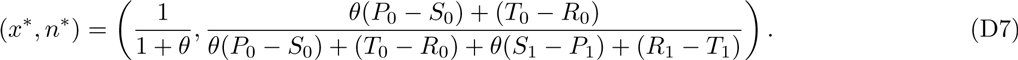

To study the stability properties, we need to compute the Jacobian matrix of *h*. We have

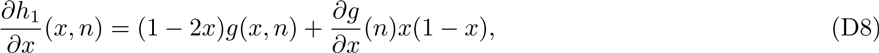

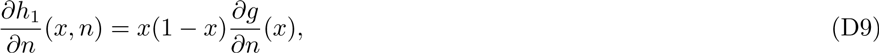

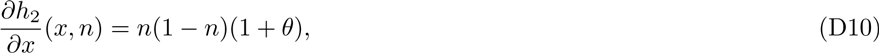

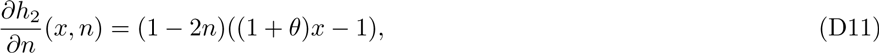

and

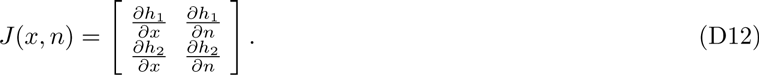

For future reference, we compute

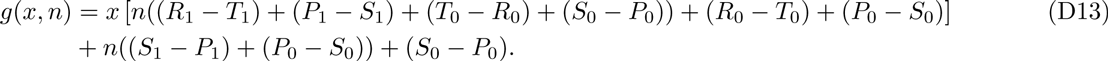

At the corner fixed points, we have

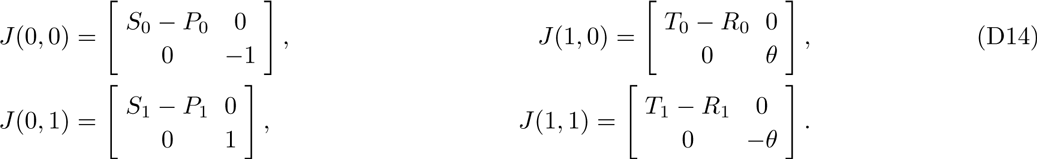

Our analysis applies to all possible combinations of conditions of the payoff matrix when *n* = 0. We denote these combinations as follows:

**Case 1 - Anti-coordination game when** *n* = 0: In the event that *R*_0_ < *T*_0_, *S*_0_ > *P*_0_.
**Case 2 - Coordination game when** *n* = 0: In the event that *R*_0_ > *T*_0_, *S*_0_ < *P*_0_.
**Case 3 - Unique Nash equilibrium by mutual cooperation when** *n* = 0: In the event that *R*_0_ > *T*_0_, *S*_0_ > *P*_0_.
**Case 4 - Unique Nash equilibrium by mutual defection when** *n* = 0: In the event that *R*_0_ < *T*_0_, *S*_0_ < *P*_0_.

Case 3 is already treated in the main text and in Section B. The possible outcomes of dynamics that do not begin at a fixed point include convergence to a stable interior fixed point or persistent oscillations in the interior. Case 4 corresponds to domination by a defector strategy when *n* = 0 and when *n* = 1. Therefore, defection will be the dominant strategy for all values of *n*. The population will converge to *x^*^* = 0 and, by extension, to the depleted environmental state *n* = 0. There are no additional dynamics possible given the feedback structure studied here. In the following sections we treat Cases 1 and 2 at length.

**FIG. S1:**
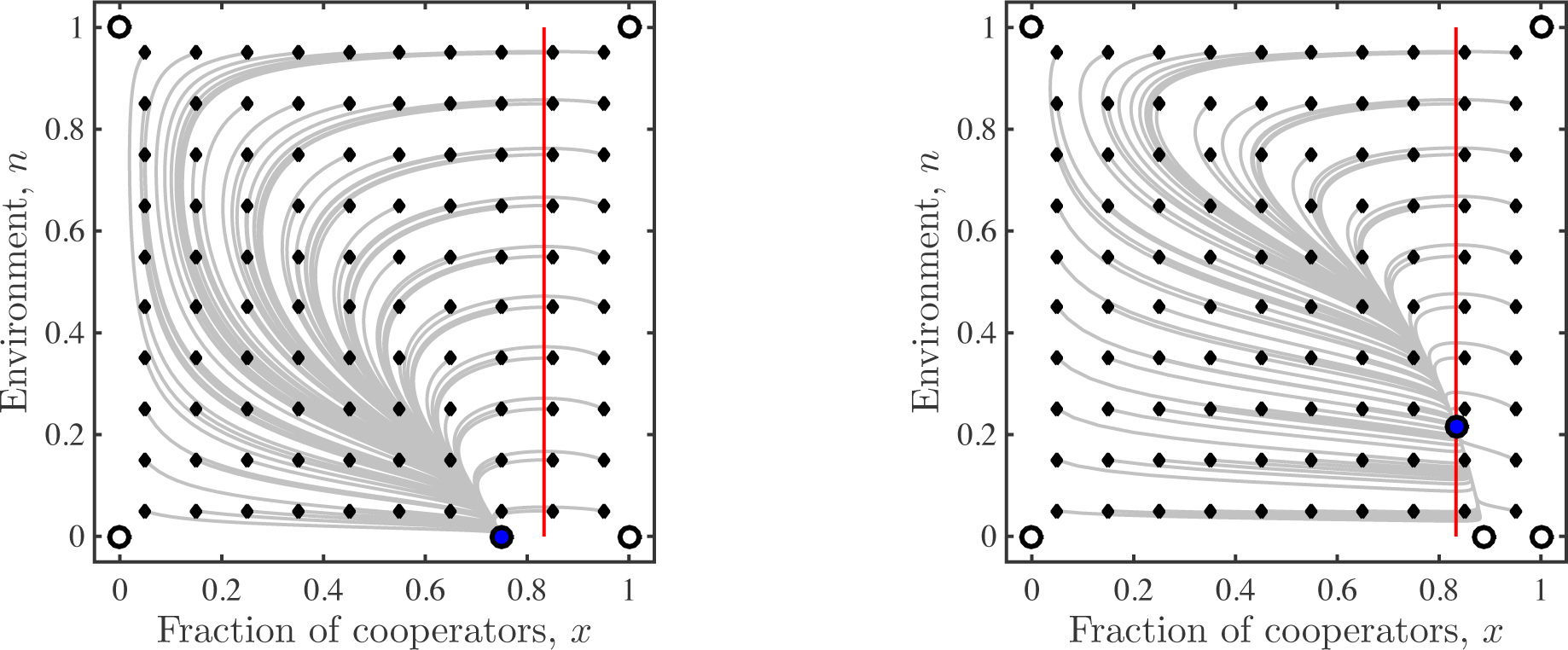
Global dynamics given an anti-coordination game in the depleted state (Case 1). The panels denote the cases when *x_m_ < x^*^* (left) and *x_m_ > x^*^* (right). The open circles denote unstable fixed points, the red line denotes the value 
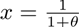
, and the solid blue circle denotes the stable fixed point. The gray lines denote trajectories initiated at the small, closed diamonds. In both cases there is a single, locally stable fixed point which is also globally stable. Numerical simulations share common parameters *θ* = 0.2 and *ϵ* = 1. (Left) Payoff matrix parameters are *R*_0_ = 5, *S*_0_ = 4, *T*_0_ = 6, *P*_0_ = 1, *R*_1_ = 3, *S*_1_ = 0, *T*_1_ = 5, and *P*_1_ = 1. (Right) Payoff matrix parameters are *R*_0_ = 5, *S*_0_ = 9, *T*_0_ = 6, *P*_0_ = 1, *R*_1_ = 3, *S*_1_ = 0, *T*_1_ = 5, and *P*_1_ = 1.

**FIG. S2:**
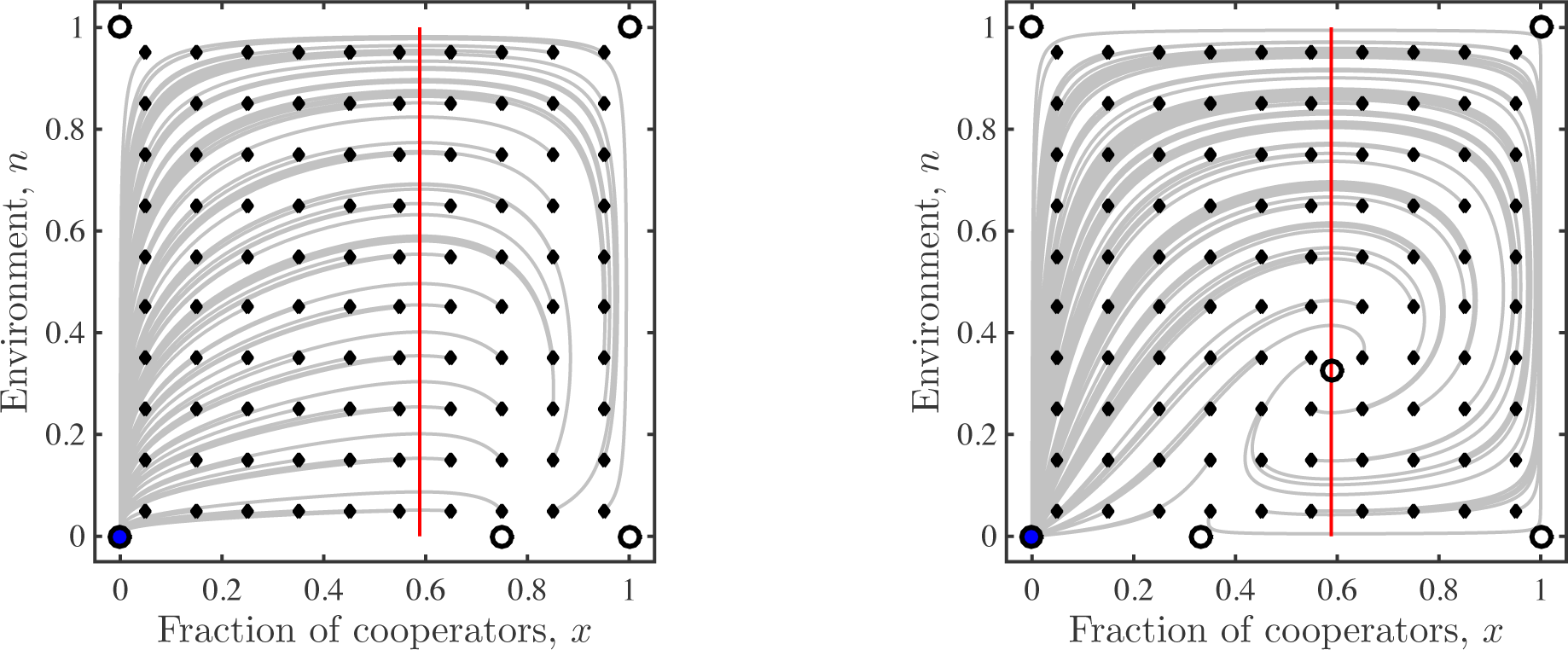
Global dynamics given a coordination game in the depleted state (Case 2). The panels denote the cases when *x_m_ > x^*^* (left) and *x_m_ < x^*^* (right). The open circles denote unstable fixed points, the red line denotes the value 
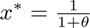
, and the solid blue circle denotes the stable fixed point. The gray lines denote trajectories initiated at the small, closed diamonds. In both cases there is a single, locally stable fixed point which is also globally stable. Numerical simulations share common parameters *θ* = 0.7 and *ϵ* = 1. (Left) Payoff matrix parameters are *R*0 = 8, *S*0 = 1, *T*0 = 7, *P*_0_ = 4, *R*_1_ = 4, *S*_1_ = 1, *T*_1_ = 5, and *P*_1_ = 2. (Right) Payoff matrix parameters are *R*_0_ = 6, *S*_0_ = 2, *T*_0_ = 4, *P*_0_ = 3, *R*_1_ = 4, *S*_1_ = 1, *T*_1_ = 6, and *P*_1_ = 2.

### Case 1 - Anti-coordination game: *R*_0_ < *T*_0_, *S*_0_ > *P*_0_

In this payoff structure, the game *A*(0) has a mixed Nash equilibrium at

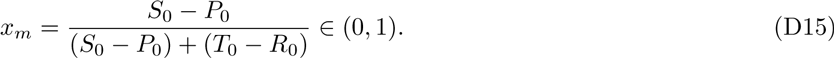

From the Jacobian analysis, all corner points are unstable fixed points. The following result characterizes the stability properties of the system.

#### Theorem D.1.

If 
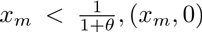
, is a stable fixed point and *n^*^* ∉ (0, 1). If 
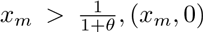
 is unstable *n^*^* ∈ (0, 1), and (*x^*^, n^*^*) is stable.

##### Proof.

We have

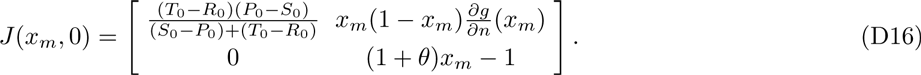

The eigenvalues *λ*_1_, *λ*_2_ are the diagonal elements of the above. We already have *λ*_1_ < 0 since *S*_0_ > *P*_0_. To ensure stability, we need (1 + *θ*)*x_m_* − 1 < 0, or when 
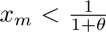
. Under this condition, *θ*(*S*_0_ − *P*_0_) < *T*_0_ − *R*_0_, which makes *n^*^* ∉ (0, 1).

For the second claim, we have 
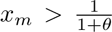
, or *θ*(*P*_0_ − *S*_0_) + (*T*_0_ − *R*_0_) < 0. This ensures that *n^*^* ∈ (0, 1), since *θ*(*S*_1_ − *P*_1_) + (*R*_1_ − *T*_1_) < 0. Also,

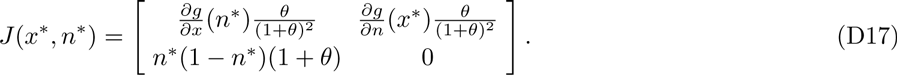

The eigenvalues are

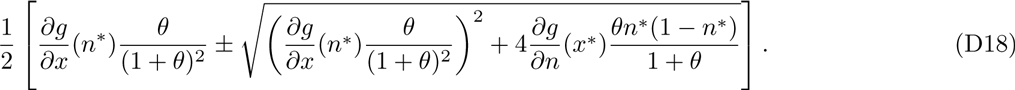

The fixed point (*x^*^, n^*^*) is stable if both eigenvalues have negative real parts. This is the case if 
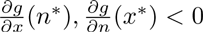
. Indeed, from (D13), we need

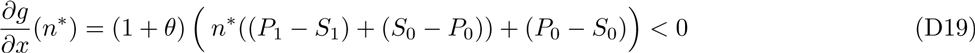

or

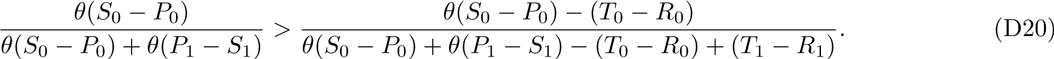

By inspection, the above is satisfied and therefore 
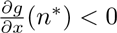
. We also need

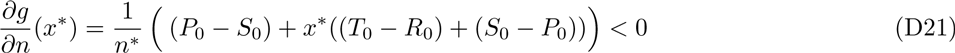

which reduces to 
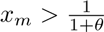
.

There are two alternative outcomes in this case. The outcomes correspond to either a single, locally stable fixed point on the *n* = 0 boundary or a single, locally stable fixed point in the interior. In both outcomes, all other fixed points are unstable. Numerical simulation confirms that these locally stable fixed points are also globally stable (see Figure S1).

### Case 2 - Coordination game: *T*_0_ < *R*_0_, *P*_0_ > *S*_0_

In this payoff structure, the game *A*(0) has a mixed Nash equilibrium at

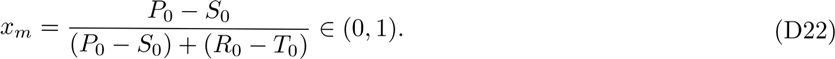

From the Jacobian analysis in (D14), the corner point (0, 0) is stable and remaining corner points are unstable. The following result characterizes the stability properties of the other (mixed) fixed points of *h*.

#### Theorem D.2.

(*x_m_*, 0) is unstable and *n^*^* ∉ (0, 1) when 
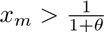
, and (*x^*^, n^*^*) is unstable when 
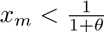
.

##### Proof.

At (*x_m_,* 0) the Jacobian have the same structure in (D16). The first diagonal element of *J*(*x*, *n*) in (D16) is positive in this case, implying *λ*_1_ > 0. Hence, (*x_m_,* 0) is unstable.

Second, we consider (*x^*^, n^*^*) with the Jacobian in (D17). First note that if 
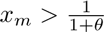
 then *n^*^* ∉ (0, 1) by (D7).

Next, we focus on the case 
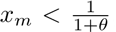
, which implies *θ*(*P*_0_ − *S*_0_) < *R*_0_ − *T*_0_. In this case, the numerator of *n^*^* is negative. The denominator of *n^*^* is a smaller negative value because *θ*(*S*_1_ − *P*_1_) + *R*_1_ − *T*_1_ < 0 hence *n^*^* ∈ (0, 1). We consider the condition 
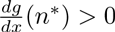
 which is given by

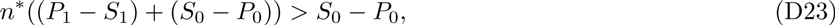

to show eigenvalues (D18) are positive. Note here that if *P*_0_ − *S*_0_ < *P*_1_ − *S*_1_ then the left hand side is positive because *n^*^* > 0. Hence the condition in (D23) is true. When the reverse holds, that is, *P*_0_ − *S*_0_ > *P*_1_ − *S*_1_, then the condition above becomes

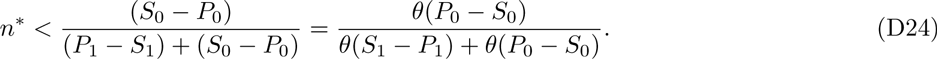

Note here that the right hand side is greater than 1 because *S*_1_ − *P*_1_ < 0. Hence the condition above holds. Therefore, 
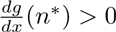
 when 
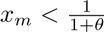
.

We conclude that there is a single, stable fixed point at (0, 0) and that all other fixed points are unstable. Numerical simulation confirms that this locally stable fixed point is also globally stable (see Figure S2).

